# A low-cost, rapidly integrated debubbler (RID) module for microfluidic cell culture applications

**DOI:** 10.1101/629642

**Authors:** Matthew J. Williams, Nicholas K. Lee, Joseph A. Mylott, Nicole Mazzola, Adeel Ahmed, Vinay V. Abhyankar

## Abstract

Microfluidic platforms use controlled fluid flow to provide physiologically relevant biochemical and biophysical cues to cultured cells in a well-defined and reproducible manner. In these systems, undisturbed flows are critical and air bubbles entering microfluidic channels can result in device delamination or cell damage. To prevent bubble entry, we report a low-cost, <u>R</u>apidly <u>I</u>ntegrated <u>D</u>ebubbler (RID) module that is simple to fabricate, inexpensive, and easily combined with existing experimental systems. We demonstrate successful removal of air bubbles spanning three orders of magnitude with a maximum removal rate (dV/dt)_max_ = 1.5 mL min^−1^, at flow rates corresponding to physiological fluid-induced wall shear stresses (WSS) needed for biophysical stimulation studies on cultured mammalian cell populations.

## Introduction

In the 1990s microfluidic systems gained popularity in analytical “lab-on-a-chip” platforms owing to unique microscale capabilities including robust sample routing, rapid multiplexed analysis, and laboratory portability [1,2]. Over the past two decades, these benefits have been extended to cell culture applications where favorable scaling effects (*e*.*g*. laminar flows, high surface to volume ratios, and short diffusion distances) have been leveraged to create physiologically-relevant microenvironments featuring precisely controlled biochemical and biophysical stimuli [3–6]. In these microscale systems, undisrupted flow is required to deliver cell culture media, maintain long-term cell viability, and control cellular-scale cues [7,8]. However, the presence of unwanted bubbles inside microscale channels can reduce media perfusion rates or create pressure buildup that lead to device failure [9]. In addition, bubbles flowing over adherent cells in culture have been shown to cause direct damage to cell membranes through exposure to dynamic air-liquid interfaces [10–12]. Given the challenges associated with unwanted bubbles in microfluidic systems, several mitigation strategies have been developed.

Bubble removal can be divided into two general approaches, i) bubble traps and ii) debubblers. Traps focus on guiding bubbles to a containment reservoir before they enter the cell culture region; traps leverage the buoyancy of air bubbles and can be vented with an external with vacuum source as the reservoir capacity is reached [13–15]. Debubblers remove bubbles via differential transport properties of liquid and air through gas-permeable membranes. For example, Xu and coworkers removed sub-microliter bubbles using a microchannel covered with a porous, hydrophobic acrylic membrane; as the flow stream made contact with the membrane, bubbles were vented through the pores in the membrane and produced a bubble-free solution downstream of the debubbler [16]. Similarly, van Lintel developed an in-line debubbler connected directly to flow tubing using a cartridge with an embedded microporous polytetrafluoroethylene (PTFE) membrane and demonstrated removal of bubbles greater than 5 µL [17]. Lui and coworkers reported a membrane-based debubbler that was integrated into a microfluidic bead array-based chip and used to prevent microliter volume bubbles from entering the detection zone of a PCR reaction [18]. These techniques have successfully removed nanoliter to microliter volume bubbles with maximum removal rates (dV/dt)_max_ ranging from 0.5 μL min^−1^ to > 2 mL min^−1^.

Although myriad debubblers have been reported in the literature, they are often complex and highly application-specific, thus integration into more general microfluidic systems represents a significant implementation barrier. To address hurdles related to both complex fabrication and integration of debubblers, we introduce a simple workflow to create a <u>R</u>apidly <u>I</u>ntegrated <u>D</u>ebubbler (RID) module that can be easily combined with existing microfluidic systems. Key practical features of RID include i) an accessible fabrication process with rapid assembly (< 2 minute), ii) low device cost (< $0.50 at lab prototype level), and iii) press-to-fit tubing connections to simplify component integration. Controlled shear stimulation of cultured cells is a hallmark capability in microfluidic systems that enables quantitative correlation between applied fluid-induced wall shear stresses (WSS) and cellular responses including endothelial cell alignment [19], calcium signaling [20], and barrier formation [21,22]. Thus, we validated RID performance by characterizing bubble removal capabilities ranging from nanoliter to microliter volume bubbles at flow rates required to apply physiological WSS to cultured mammalian cells within standard geometry microfluidic channels.

## Materials and Methods

### RID fabrication and assembly

As shown in **Figure 1**, structural elements (L1-L3) were designed as vector files in Adobe Illustrator and cut from PMMA (2 mm thickness, McMaster-Carr) using a 40W CO_2_ laser (Full Spectrum Laser, H-Series) with inlet and outlet ports in L1 designed to house #003 rubber O-rings (OD 3/16”, ID 1/16”, Durometer 70A, McMaster-Carr). Pressure sensitive adhesive films (PSA, 3M MP467) were rolled onto the top surfaces of L1-L3 prior to laser cutting using a cold roll laminator. A polytetrafluoroethylene (PTFE) membrane (Sterilitech, 0.01 mm pore diameter, 0.1 mm thickness) was cut with a razor blade and placed in contact with the PSA on L3. L2 (PMMA) was used to protect the PTFE membrane from damage due to improper insertion of the tubing. The 2 mm wide by 12 mm region cut into L5 (polyester film) defined the degassing region of the device, and the PSA lamination produced liquid-tight sealing between layers. Once all layers were cut in a batch process, individual devices could be assembled in less than 2 minutes. The packed O-ring assembly allowed simple press-to-fit connection with 1/16” OD tubing and the addition of barbed end fittings (Cole Parmer) allowed attachment to flexible tubing. The overall footprint of the RID module was 10 mm x 18.5 mm (W × L).

**Figure 1.**
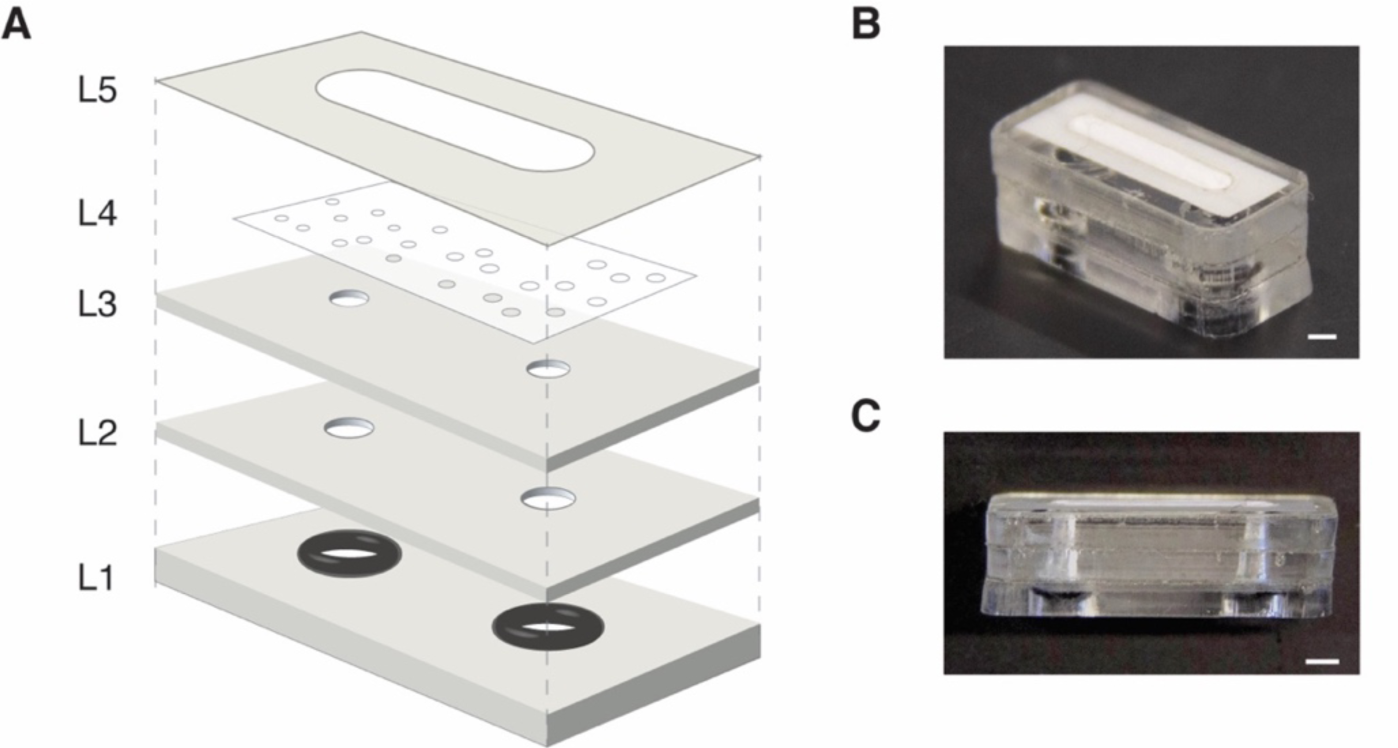
A) Exploded view of RID. L1 – L3 are laser cut from PMMA. L1 and L2 have pressure sensitive adhesive on the top face to facilitate layer-to-layer lamination. L3 is the tubing stop layer used to prevent damage to L4, the porous, hydrophobic PTFE membrane. L5 is laser cut from polyester film and is used to seal the edges of the PTFE membrane to prevent leakage. After assembly, an O-ring is inserted into the access ports in L1 to provide simple push-to-connect compression sealing to interface the device with tubing for fluid routing. B) Isometric image of assembled RID module. Scale bar = 2 mm. C) Side view of assembled device. Scale bar = 2mm.

### RID operating principle

When the pressure from the input fluid stream, P, exceeded the opening pressure, P_open_, L4 deflected away from L3 and exposed a fluidic path from the inlet to the outlet. Air bubbles present in the flow stream made contact with the PTFE membrane and were vented out of the device due to the transmembrane pressure gradient between the segmented flow and the atmospheric pressure, P_atm_. However, liquid segments were unable to move through the hydrophobic membrane at pressures less than the critical membrane liquid entry pressure, P_critical_, and recombined into to form a single-phase flow prior to exiting RID. The operating pressure range was therefore P_open_ < P < P_critical_.

### RID characterization and automated image analysis

To characterize RID bubble removal, individual streams of air and liquid (New Era Pump Systems, Inc) were combined to create segmented flow streams and directed toward the input port of RID. The test liquid was complete cell culture media (DMEM/F12, 10% v/v FBS) with food color added increase contrast during imaging. Flow streams entering and exiting the RID module were simultaneously recorded (Samsung Galaxy S6) and analyzed with custom a MATLAB image processing script (available upon request). Black and white binary images corresponding to air and liquid respectively were obtained from the captured video frames. The leading and trailing edges of the segments were identified, and tubing inner diameter along with video frame rate were used to calculate i) the flow rate Q, ii) volume of each bubble V_b_, and iii) V_b_:V_T_ ratio (defined as bubble volume to total volume). A test condition failed when a bubble was visualized in the fluid exiting the device via image analysis. We conservatively report the failure threshold as the lower bound of the 95% confidence interval determined from failure testing of four independent RID devices. The equation WSS = 6μQw^−1^h^−2^ (valid for laminar, fully developed flow and h << w) [23], was used to relate Q to applied wall shear stress (for a bubble-free stream) in a standard geometry microfluidic culture channel with h = 0.1 mm, w = 1 mm, l = 10 mm (channel volume, V_channel_ = 1 μL).

### RID opening pressure P_open_, and membrane liquid entry pressure P_critical_

A 5mL fluid reservior was connected vertically to the input of the debubbler. With the debubbler dry, cell culture media was added to the reservoir to generate a hydrostatic pressure head ΔP = ρgΔh with Δh denoting the difference in height between the media level and the outlet port. Using video analysis, the height difference required to initiate flow through the debubbler was determined and converted to the opening pressure P_open_. Results reported as mean ± standard deviation (SD) from four independent devices.

A syringe pump was used to introduce cell culture medium to RID while the outlet port was connected to dead-end tubing. Pressure sensors (Parker Hannifin) on either side of the debubbler measured the pressure across the module as a function of time. Video analysis was used to determine the critical liquid pressure P_critical_ at which fluid was forced through the PTFE membrane (see Supplemental). Results reported as mean ± SD from four independent devices.

### Fluid properties, contact angle, and interfacial tension measurements

A glass capillary viscometer was used to measure the viscosity of DMEM F12 cell culture media with 10% FBS. Fluid density was determined by measuring mass of a known media volume using an analytical balance (Mettler Toledo) at 37°C. A goniometer (Rame-Hart 250) was used to measure the interfacial tension of cell culture media in air, and the contact angle between the culture medium and PTFE membrane using the hanging drop and sessile drop methods respectively. Results reported as mean ± SD from four independent devices.

### Cell Culture

For routine culture, human umbilical vein endothelial cells (HUVEC) were cultured in DMEM-F12 with 10% v/v fetal bovine serum using manufacturer’s protocols. Briefly, HUVECs were enzymatically disassociated using Trypsin/EDTA (Life Technologies) and sub-cultured when they reached 70% confluence. Cells were centrifuged at 250g, resuspended in medium, and counted. Cells were plated at 5,000 – 10,000 cells cm^−2^ on Geltrex coated tissue culture flasks. Media was replaced every 24-48 hours.

For bubble exposure studies, a microfluidic channel network was sealed against a glass slide using our previously reported reversibly <u>S</u>ealed <u>E</u>asy <u>A</u>ccess <u>M</u>odular (SEAM) platform [24,25]. Glass slides (Fisher) were sterilized in ethanol, rinsed three times in PBS, then allowed to dry in a biosafety cabinet. A PMMA piece with a 15 × 20 mm cavity and embedded magnets was sterilized with ethanol attached to the glass with PSA. The cavity was treated with a Geltrex solution (Thermo Fisher, 0.1 mg mL^−1^) at 37°C for one hour to improve cell adhesion. Excess Geltrex solution was aspirated, and a PDMS microfluidic network was magnetically sealed against the glass substrate. The microfluidic channel network was fabricated using a soft lithography process described previously [24]. HUVECs were seeded at a density of 40,000 cells cm^−2^ and allowed to attach overnight. Cells were then treated with Calcein AM (ThermoFisher, 0.5 μM concentration in PBS) and the cell permeable Hoechst 33342 stain (ThermoFisher, 1:2000 dilution in PBS) for 15 minutes to aid visualization of HUVEC cytoplasm and nuclei respectively. HUVECs in culture were imaged after exposure to a segmented air-liquid stream with flow rate corresponding to WSS of 11.5 dyne cm^−2^ with and without RID module upstream of the culture device. RID modules were sterilized with ethanol followed by 3X wash with PBS to remove residual ethanol, and allowed to dry before use.

## Results and Discussion

### Characteriation of RID bubble removal capabilities

To ensure that RID could be used for shear stimulation studies relevant to human (1-50 dyne cm^−2^) [26] and rodent cell studies (50-200 dyne cm^−2^) [27], segmented air-liquid streams were introduced at flow rates corresponding to defined WSS in a standard geometry microfluidic channel, h = 0.1 mm, w = 1 mm, l = 1 cm, with the channel volume V_channel_ = 1 μL. The range of tested bubble volumes was selected to evaluate RID performance under common flow disruption scenarios: i) V_b_ < V_channel_, ii) V_b_ ≥ V_channel_ iii) V_b_ >> V_channel_ (*i*.*e*. catastrophic problem with pump or pressure source). A custom image processing algorithm was used to measure V_B_, V_b_:V_T_, and Q. Results from RID bubble removal testing are summarized in **Figure 3** with A test condition was considered to be unsuccessful if a single bubble was detected in the outlet tubing via image analysis; therefore, the reported operational range represents conditions where air bubbles were completely removed.

**Figure 2.**
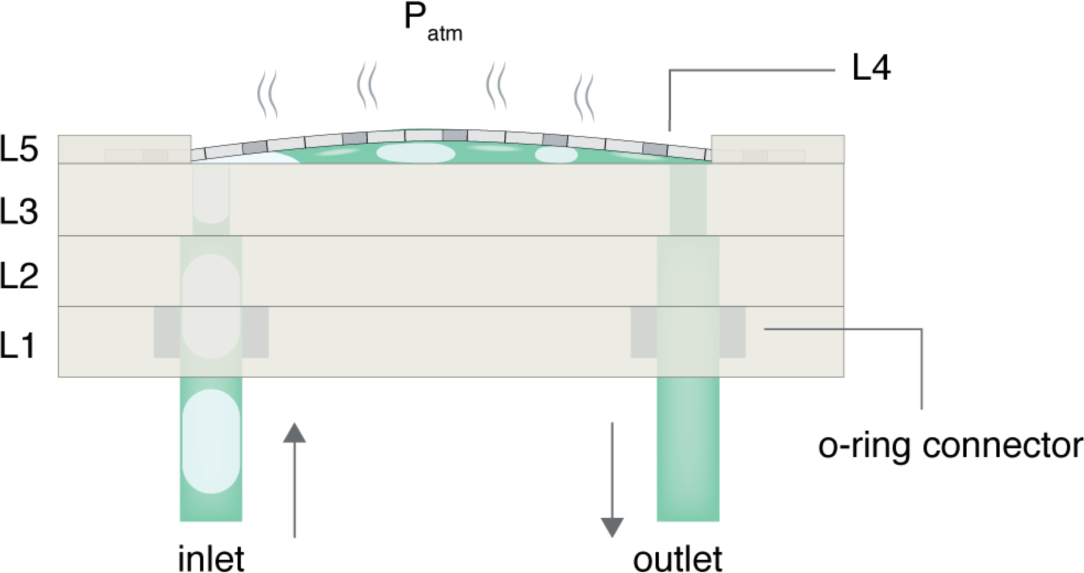
Schematic representation of RID in operation. Once the opening pressure P_open_ is exceeded, the PTFE membrane (L4) deflects and exposes a fluidic path connecting the inlet and outlet. Bubbles in the input stream make contact with the hydrophobic PTFE membrane and are removed as a result of the differential pressure between the air-liquid fluid stream and the atmosphere. The liquid is unable to move through the membrane below the critical liquid entry pressure, P_critical_. As air bubbles are removed, liquid plugs recombine to form a single-phase flow exiting RID.

**Figure 3.**
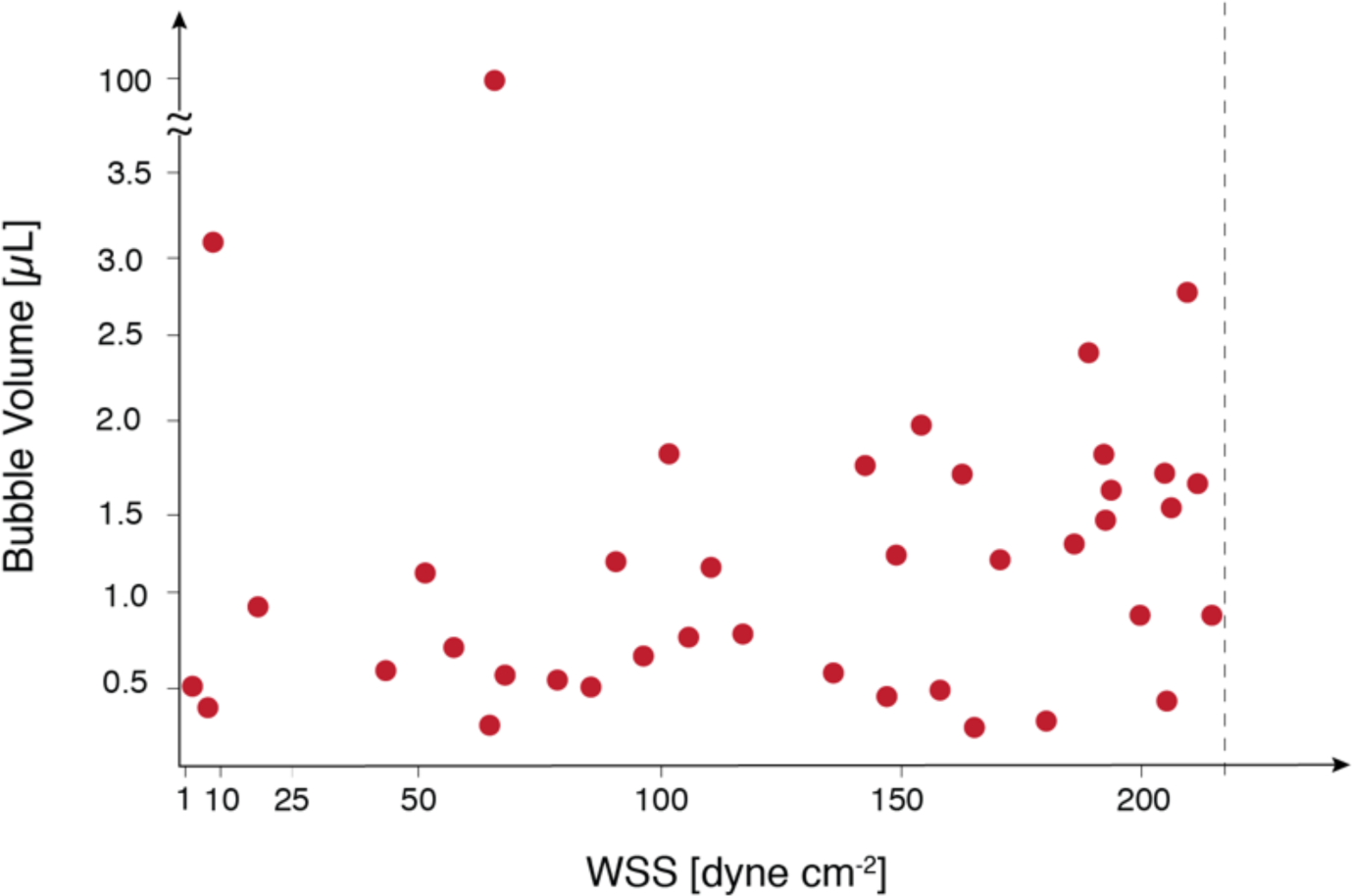
RID air bubble removal results at flow rates corresponding to defined WSS in a microfluidic channel. Each data point represents air bubbles successfully removed from a segmented air-liquid flow with bubble volume V_b_. The dashed line represents the lower bound of the 95% confidence interval for successful device operation (WSS = 213 dyne cm^−2^, Q = 2.6 mL min^−1^). Air bubbles ranging from 200 nL to 100 μL were successful removed and the maximum air removal rate (dV/dt)_max_ = 1.5 mL min^−1^. RID is operational across the physiological WSS range needed for *in vitro* shear stimulation studies using both human and rodent cells.

RID successfully removed bubbles across three orders of magnitude (200 nL ≤ V_b_ ≤ 100 μL) from segmented flows (0.07 ≤ V_b_:V_T_ ≤ 0.7) including the challenging high Q (high WSS), small V_b_, high V_b_:V_T_ conditions where dV/dt must be sufficiently high to remove bubbles before they were displaced downstream and out of the device by the incoming flow. The average WSS at failure = 289 ± 48 dyne cm^−2^ with a 95% confidence interval (213 < WSS < 352 dyne cm^−2^); the dashed line in **Figure 3** shows the lower limit of the 95% confidence interval (213 dyne cm^−2^ or Q = 2.6 mL min^−1^), and represents a conservative limit of the operational range. Although the 95% confidence interval is relatively large, RID reproducibly removed large and small volume bubbles at flow rates corresponding to WSS that span the physiological ranges of both human and rodent cells that are commonly used to study shear stimulation *in vitro*. RID (dV/dt)_max_ = 1.5 mL min^−1^ and enables rapid removal of bubbles and is competitive with more complex and specialized debubblers. Traditionally, debubblers are designed for a particular bubble removal operation range; to the best of our knowledge, this is the first demonstration of a rapid debubbler spanning at least three orders of magnitude in removed V_b_ with a single device. It should be noted that V_b_ = 100 μL was the maximum bubble volume tested due to experimental limitations, and was not the operational failure limit of RID. Although bubbles with V_b_ < 200 nL could not be reproducibly generated using our experimental setup, we anticipate that smaller volume bubbles would be removed either at i) the entrance of RID via contact with the PTFE membrane due to the perpendicular orientation of the input tubing or ii) in the gap between the membrane and PMMA where the bubble is forced to make contact with the membrane.

**Figure 4** shows a representative image sequence spanning two seconds and demonstrating removal of air bubbles (V_b_ = 2.2 ± 0.3 μL, V_b_:V_T_ = 0.4) at Q = 1.1 mL min^−1^ (liquid WSS = 90 dyne cm^−2^); flow direction is left to right. Bubbles entering the RID module were removed with dV/dt = 440 μL min^−1^ and single-phase flow exited the device. See supplemental information for full video.

**Figure 4.**
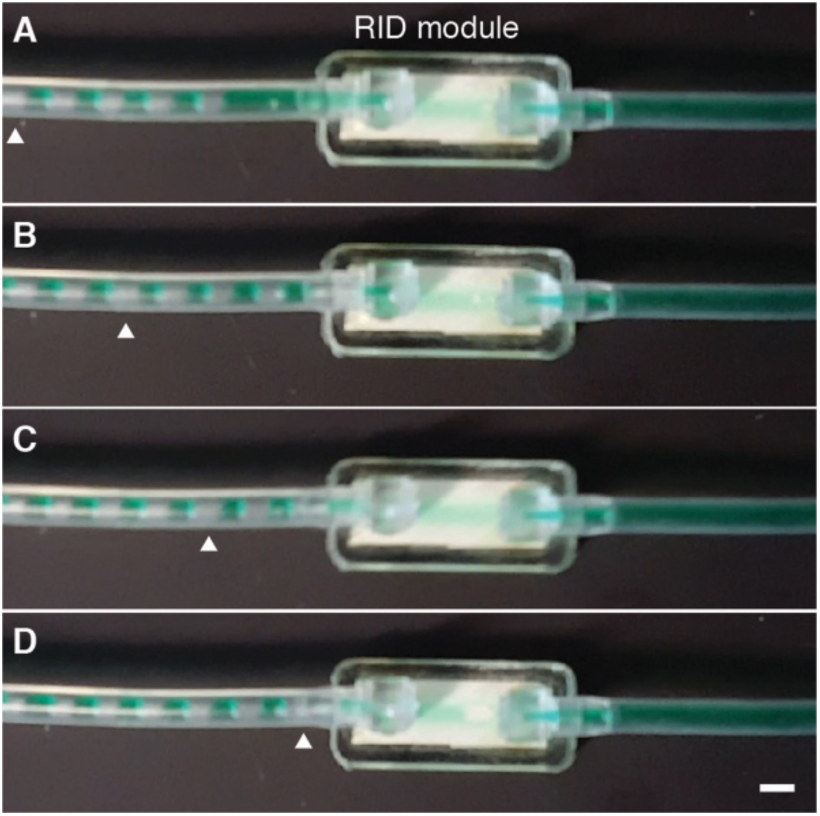
Sequential frames A-D from extracted from video showing bubble removal from an incoming segmented flow stream (bubble volume = 2.2 ± 0.3 μL, Q = 1.1 mL min^−1^ and V_b_:V_T_ = 0.4). Flow is left to right. A) White arrow highlighting bubble at left edge of frame, t = 0 sec. B) Segmented flow moving down tube toward RID, t = 0.73 sec. C) Bubbles upstream of highlighted bubble enter RID, t = 1.33 sec. D) Highlighted bubble close to RID entrance. Upstream bubbles have been removed bubble-free flow leaves RID, t = 2 sec. Bubble removal rate dV/dt = 440 μL min^−1^. Scale bar = 3mm.

### 3.2 Cell damage resulting from bubble introduction

As shown in **Figure 5**, we demonstrate the importance of preventing air bubbles from entering a microfluidic channel. To replicate a common flow disturbance where V_b_ ≥ V_channel_, segmented flow streams were introduced upstream of the culture module with and without RID in place. Flow rate corresponding to physiological WSS of 11.5 dyne cm^−2^ (for bubble-free flow) was maintained for both experimental conditions. **Figure 5** A) and C) schematically show the experimental conditions (e.g. segmented flows +RID and −RID), while B) shows viable cells in the culture channel with RID in place. D) Without RID in place membranes appear damaged. The damage can be explained in part by an amplification in applied WSS resulting from a thin lubrication layer present between the bubble and surrounding walls [11,28,29]. WSS amplification is caused by an increased velocity gradient in the thin lubricating layer of thickness b << h as the bubble moves through the channel, compared to the bubble-free condition where WSS = 6μQw^−1^h^−2^. Under the test conditions, we estimate the amplification factor Ф comparing WSS with and without bubbles (e.g. WSS_bubble_/WSS) = 55 (see supplemental for details). Thus, the presence of bubbles in a microfluidic channel could dramatically increase WSS from physiological conditions to levels where cell damage can occur. Introduction of RID in the flow path prevented bubbles from entering the channel network and supported viable cell culture. RID fabrication workflow enables complete debubbler customization to meet experimental needs and simplify integration into a microfluidic experimental setup.

**Figure 5.**
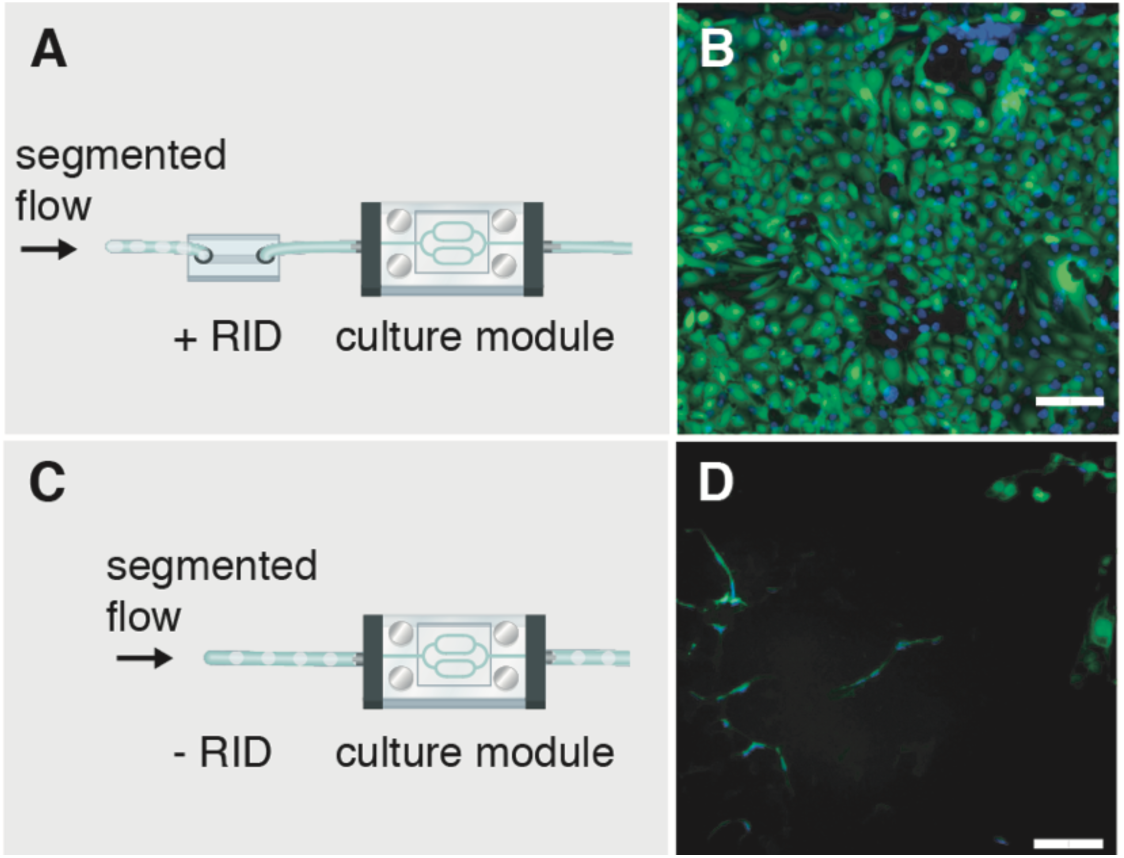
Representative images showing the importance of RID to protect against bubble-induced damage. Segmented flows were introduced upstream of the cultured cells A) with RID and C) without RID in place. B) With RID in place, the cell population was viable while D) without RID cells bubbles entering the channel caused cell damage. Scale bar = 100 μm.

### RID operating pressure range and chip-to-world interconnections

To determine the operating pressure range for RID (P_open_ < P < P_critical_) we measured i) the minimum pressure, P_open_, required to deflect the PTFE membrane from the PMMA layer and ii) the maximum pressure, P_critical_, above which cell culture media was forced through the hydrophobic PTFE membrane pores (see supplemental figure). The operating pressure range spanned three orders of magnitude with P_open_ = 7.4 ± 0.4 Pa and P_critical_ = 9.7 ± 1.7 kPa. The small opening pressure can be attributed to the limited adhesion between the hydrophobic PTFE and the PMMA surface that must be overcome by the flow stream pressure. P_critical_ is a function of membrane pore diameter; the manufacturer’s reported PTFE pore diameter of 10 um was sufficient for our applications. However, a membrane with a smaller pore size can be used if a higher P_critical_ is desired. It should be noted that Sterilitech PTFE membranes do not contain well-defined circular pores but are comprised of a fibrous mesh that contain voids between fibers; the relatively large 95% CI for WSS at failure can be attributed in part to inhomogeneities between membranes.

Microfluidic cell culture platforms often require elements to be connected together to form a fluidic flow path (e.g. a pressure source sequentially connected to a flow damper, cell culture module, and downstream collection vessel). The interconnection problem can be a source of frustration because each component often has different tubing requirements and mating mechanisms (e.g. barbed, press to fit, or ferrule). A goal of this work is to increase accessibility by providing an integration approach to simplify connections. Here, we implemented a simple connection mechanism via embedded O-rings in the top housing layer of the RID module that form a compression seal against inserted rigid tubing. The press-to-fit O-ring connector (See Figure 2) was able to withstand pressures of at least 70kPa (maximum range of pressure sensor), which was sufficient to ensure leak-proof operation in our system where the maximum operatonal pressure Pcritical is an order of magniture lower than 70kPa. The O-ring connector can also be used as an adaptor to insert barbed fittings to connect RID with flexible tubing and thus simplfy integration into existing microfluidic setups without requiring design modification.

### Device fabrication workflow

To facilitate customization and improve debubbler accessibility for the microfluidics community, the RID fabrication workflow uses equipment commonly found in a community makerspace or general research laboratory. RID incorporates commercially available materials (e.g. PMMA, PSA, and PTFE membranes) and modules can be assembled in less than 2 minutes when layers (Figure 1) are cut in a batch process. Design prototypes can be easily iterated because the process workflow allows complete control over the device geometry and layer components. With the current design, each module costs less than $0.50 (see Table S1), allowing them to be treated as consumable components. The fabrication process is also amenable to scale up using reel-to-reel processing or injection molding techniques once a final design is identified.

## Conclusions

We have demonstrated a simple fabrication workflow to create a debubbler that can rapidly remove bubble volumes spanning three orders of magnitude from segmented flows at flow rates compatible with those required for microfluidic shear stimulation studies. The fabrication process can be carried out in a general makerspace or research laboratory and the footprint and fluidic interconnections can be customized as needed to fit existing experimental setups. We anticipate that the combination of simple fabrication, integration, and functional capabilities will enable RID to be easily implemented into microfluidic applications where the entry of bubbles is undesired.

## Supporting information

supplemental equation and table

supplemental video – time sequence

supplemental video – RID operation

## Supplementary Materials

Supplemental information: WSS amplification calculation and RID Cost. Video S1: Rapid bubble removal. Video S2: Figure 4 Video.

## Author Contributions

This project was conceptualized by V.V.A. and M.J.W. MATLAB code was developed by M.J.W. and N.K.L. Formal analysis and data acquisition was performed by all authors. All authors contributed to draft preparation, review, and editing. Project was supervised by V.V.A.

## Funding

This work was supported by the Kate Gleason College of Engineering New Faculty Startup Fund

## Acknowledgments

The authors acknowledge Cristian Almendariz and Mohammad Raziul Hasan for their early stage contributions to this work and thank the Schertzer Lab at RIT for assistance with contact angle and interfacial surface tension measurements. Illustration credit to Brad Kwarta.

## Conflicts of Interest

The authors declare no conflict of interest. The funders had no role in the design of the study; in the collection, analyses, or interpretation of data; in the writing of the manuscript, or in the decision to publish the results.

