## supplemental equation and table for "A low-cost, rapidly integrated debubbler (RID) module for microfluidic cell culture applications"

**Supplemental Information**

*Amplification of fluid-induced wall shear stress with bubble introduction*

Applied WSS in a rectangular channel the absence of bubbles can be determined using the relation WSS = $6\mu Q w^{-1} h^{-2}$ for laminar, fully developed flow. In this experiment, Q corresponded to 11.5 dyne $cm^{-2}$ in a microfluidic channel with h = 0.1 mm, w = 1 mm, l = 10 mm. As a first order estimate of WSS amplification in the presence of well-studied Taylor bubbles, (e.g. bubble length > channel width) it is assume a lubrication film is present between the bubble and the surrounding walls [1,2]. In the classical analysis, film thickness is considered to be constant between the wall and the Laplace pressure jump across the curved air-liquid interface at the front of the bubble is neglected. The capillary number Ca = $\mu U \gamma^{-1}$ with bubble velocity U, and interfacial tension $\gamma$ and relates viscous to interfacial phenomena; for Ca < $5x10^{-3}$, the Bretherton equation can be used to estimate the thickness of the lubrication film: b = $0.643(3\ Ca^{2/3})R_H$ where $R_H$ is the hydraulic radius. For channels with w >> h, $R_H$ simplifies to 0.5h. In our system, $\mu$ =$8.24x10^{-4}$ Pa sec, $\gamma$ = 0.063 N $m^{-1}$, (both measured at 37°C) and bubble velocity U measured for input flow rate of Q = 140 µL $min^{-1}$, Ca = $3.06x10^{-4}$ and film thickness b = 0.3 µm. With bubble velocity U, $WSS_{bubble}$ = $\mu U b^{-1}$ and amplification factor $\Phi = WSS_{bubble}/WSS$. In our system, $\Phi$ = 55 and is consistent with similar analysis by Lochovsky and coworkers [3].

**Table 1 – List of Materials and Estimated Cost Per Device**

| Material | Cost/$mm^2$ [USD] | Total cost per device | Layer designation |
|---|---|---|---|
| PMMA | 0.0000569 | $0.0316 | L1-L3 |
| PSA | 0.0000503 | $0.0279 | L1-L3 |
| PTFE | 0.002334 | $0.2987 | L4 |
| O-rings | 0.034 (per O-ring) | $0.0680 | L1 |
| transparency | 0.0000325 | $0.0060 | L5 |
| | | **$0.43** | |

1. Bretherton, F.P. The motion of long bubbles in tubes. J. Fluid Mech. 1961, 10, 166.
2. Klaseboer, E.; Gupta, R.; Manica, R. An extended Bretherton model for long Taylor bubbles at moderate capillary numbers. Phys. Fluids 2014, 26, 32107.
3. Lochovsky, C.; Yasotharan, S.; Günther, A. Bubbles no more: In-plane trapping and removal of bubbles in microfluidic devices. Lab Chip 2012, 12, 595–601.
